# Structural basis for inhibition of mycobacterial ATP synthase by squaramides and second generation diarylquinolines

**DOI:** 10.1101/2023.02.03.527018

**Authors:** Gautier M. Courbon, Paul R. Palme, Lea Mann, Adrian Richter, Peter Imming, John L. Rubinstein

## Abstract

Mycobacteria, such as *Mycobacterium tuberculosis*, depend on the activity of adenosine triphosphate (ATP) synthase for growth. The diarylquinoline bedaquiline (BDQ), a mycobacterial ATP synthase inhibitor, is an important medication for treatment of drug-resistant tuberculosis but suffers from off-target effects and is susceptible to resistance mutations. Consequently, both new and improved mycobacterial ATP synthase inhibitors are needed. We used electron cryomicroscopy and biochemical assays to study the interaction of *Mycobacterium smegmatis* ATP synthase with the second generation diarylquinoline TBAJ-876 and the squaramide inhibitor SQ31f. The aryl groups of TBAJ-876 improve binding compared to BDQ, while SQ31f, which blocks ATP synthesis ~10 times more potently than ATP hydrolysis, binds a previously unknown site in the enzyme’s proton-conducting channel. Remarkably, BDQ, TBAJ-876, and SQ31f all induce similar conformational changes in ATP synthase, suggesting the resulting conformation is particularly suited for drug binding. Further, high concentrations of the diarylquinolines uncouple the transmembrane proton motive force while for SQ31f they do not, which may explain why high concentrations of diarylquinolines have been reported to kill mycobacteria while SQ31f has not.

## Introduction

Tuberculosis (TB) is the second most common cause of death by infectious disease worldwide after COVID-19 (World Health Organization, 2022). Infection by *Mycobacterium tuberculosis*, which causes TB, has started to develop resistance to first-line antibiotics (Murray et al., 2022), jeopardizing efforts to eradicate the disease and necessitating development of new antibiotics to treat drug resistant TB. The recently-approved diarylquinoline drug bedaquiline (BDQ, Fig. 1A, *blue*) targets mycobacterial adenosine triphosphate (ATP) synthase (Andries et al., 2005) and has been added to WHO’s list of essential medicines for the treatment for multidrug-resistant TB (World Health Organization, 2019). BDQ kills *M. tuberculosis* because of the organism’s reliance on aerobic respiration (Cook et al., 2017). During respiration, the electron transport chain pumps protons from the cytosol across the plasma membrane, establishing a transmembrane proton motive force (PMF). This PMF drives protons back into the cytosol through a periplasmic half channel and a cytosolic half channel in the ATP synthase’s membrane-embedded F_O_ region, inducing rotation of a rotor subcomplex that drives ATP synthesis in the enzyme’s catalytic F_1_ region (Courbon and Rubinstein, 2022). BDQ binds five low-affinity sites involving the ring-forming c subunits from the membrane-embedded portion of the rotor (Koul et al., 2007; Preiss et al., 2015) as well as two high-affinity sites that also involve the a subunit that forms the proton conducting pore across the membrane (Guo et al., 2021). BDQ binding blocks rotation of the rotor, inhibiting ATP synthesis at nanomolar concentrations, leading to bacteriostasis with a minimum concentration for inhibition of 99% of growth (MIC_99_) of 0.03 μg/mL (54 nM) for *M. tuberculosis* H37Rv (Andries et al., 2005). However, at micromolar concentrations, BDQ also allows protons to translocate across the mycobacterial inner membrane without synthesizing ATP, thereby dissipating the PMF (Hards et al., 2015, 2018). Maintaining a PMF is essential for *M. tuberculosis* viability and consequently this uncoupling activity has been proposed to be important for the killing of mycobacteria that occurs with prolonged exposure to micromolar concentrations of BDQ (> 30 × MIC_99_) (Koul et al., 2014; Hards et al., 2015; Rao et al., 2008). Consistent with the observed uncoupling, nanomolar concentrations of BDQ inhibit ATP hydrolysis by detergent-solubilized ATP synthase but micromolar concentrations of BDQ restore ATP hydrolysis activity (Guo et al., 2021). However, it is unclear if the micromolar restoration of ATP hydrolysis by the detergent-solubilized enzyme and the micromolar uncoupling of the PMF when ATP synthase is in a membrane are both caused by disrupting the enzyme structure (Feng et al., 2015; Hards et al., 2018).

**Figure 1.**
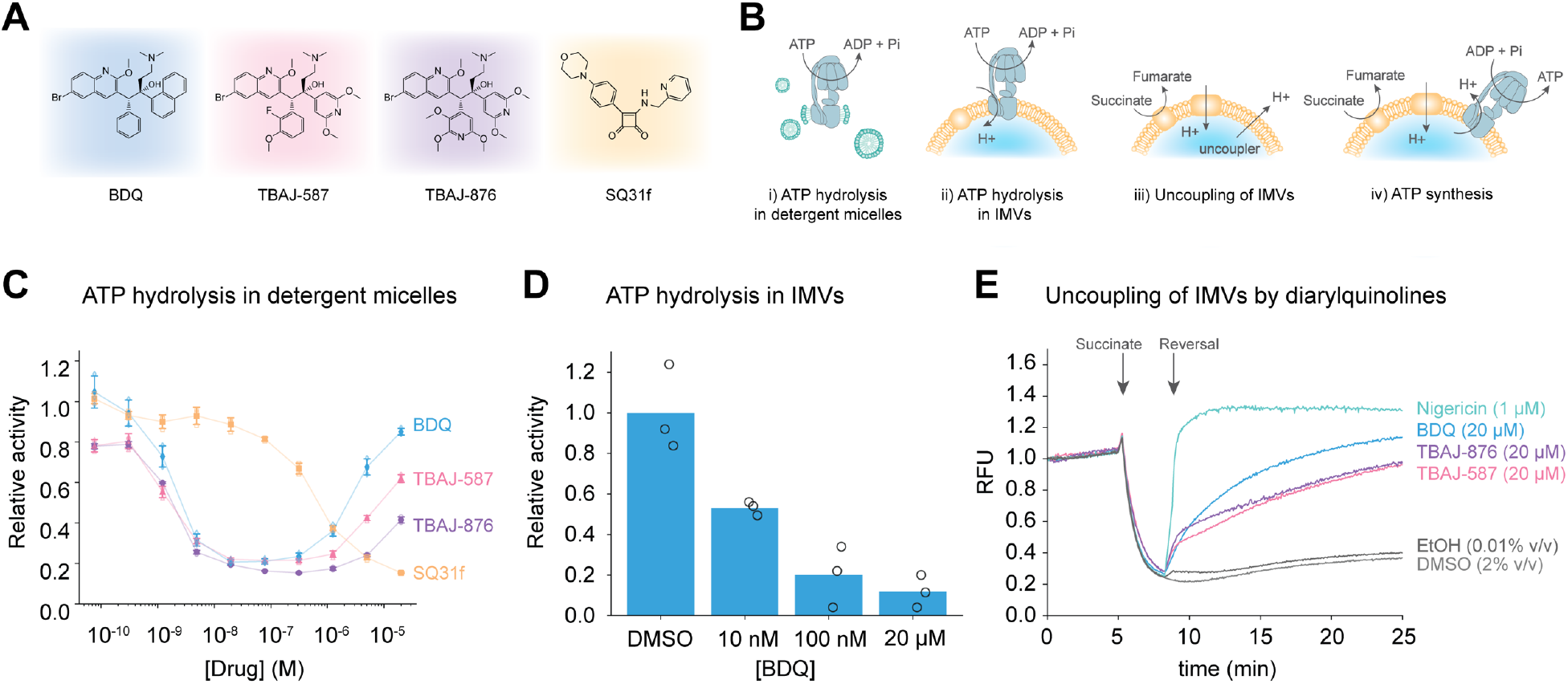
ATP hydrolysis inhibition and IMV uncoupling. **A,** Chemical structure of BDQ, TBAJ-587, TBAJ-876, and SQ31f. **B,** Schematic representation of the assays performed: ATP hydrolysis in detergent micelles (i), ATP hydrolysis in IMVs (ii), uncoupling of IMVs (iii), and ATP synthesis by IMVs (iv). **C,** Diarylquinoline and SQ31f inhibition of ATP hydrolysis by purified ATP synthase bearing truncated α subunits. Mean relative activity (filled symbols) ± s.d from n=3 separate assays (empty symbols) for each of four different inhibitors (triangles, circles, diamonds, and squares). BDQ reference curve is from (Guo et al., 2021). **D,** Inhibition of ATP hydrolysis activity in *M. smegmatis* IMVs containing ATP synthase with truncated α subunits. Initial ATP hydrolysis activity of the ATP synthase is defined as the activity that is sensitive to 100 μM DCCD. Mean relative activity from n=3 separate assays is shown. **E**, Uncoupling activity of diarylquinoline analogs in *M. smegmatis* IMVs monitored with ACMA fluorescence. Each compound was assayed separately, and the initial fluorescence was set to 1.0. RFU: relative fluorescence units.

Despite its success, lipophilicity and off-target activities of BDQ can cause side effects such as hepatotoxicity and cardiac arrythmia (Mase et al., 2013). Indeed, the high calculated log octanol-water partition coefficient of BDQ (cLogP=7.25) and its relatively low half-maximal inhibitory concentration of the cardiac potassium channel hERG (IC_50_=1.6 μM) lead to accumulation of the drug in tissues and can prolong the QT interval of the heart, respectively (Sutherland et al., 2019). In an effort to reduce BDQ lipophilicity and hERG channel inhibition the second generation diarylquinolines TBAJ-587 (Fig. 1A, *pink*) and TBAJ-876 (Fig. 1A, *purple*) were developed (Sutherland et al., 2019). These compounds differ from BDQ due to changes in the two aryl groups while keeping the rest of the scaffold untouched. In TBAJ-587, BDQ’s phenyl group is replaced with a 2-fluoro-3-methoxy phenyl moiety and its naphthalenyl group is replaced with 2,6-dimethoxy pyridine. These changes decreased cLogP to 5.8 and abate hERG inhibition to IC_50_=13 μM while also improving the MIC_90_ against *M. tuberculosis* from 0.04 to 0.006 μg/mL (Sutherland et al., 2019). In TBAJ-876, BDQ’s phenyl group is replaced with 2,3,6-trimethoxy pyridine while its naphthalenyl group is replaced with 2,6-dimethoxy pyridine. These modifications further improved solubility to clogP=5.15 and maintain a low MIC_90_ of 0.004 μg/mL while drastically reducing inhibition of the hERG channel to IC_50_>30 μM (Sutherland et al., 2019). The efficacy of TBAJ-876 and TBAJ-587 have been demonstrated both *in vitro* and *in vivo* in murine models of TB (Sutherland et al., 2019; Xu et al., 2021) and the compounds are currently undergoing phase I clinical trials (https://clinicaltrials.gov/ct2/show/NCT04493671 and https://clinicaltrials.gov/ct2/show/NCT04890535).

As clinical use of BDQ increases, BDQ-resistant *M. tuberculosis* strains have begun to emerge. The leading cause of resistance to BDQ is upregulation of MmpL5/MmpS5-mediated drug efflux, which results in a moderate increase in the drug’s MIC (Villellas et al., 2017). Resistance mutations in the mycobacterial ATP synthase remain rare clinically (Huitric et al., 2010; Zimenkov et al., 2017; Andres et al., 2020) but these mutations can lead to complete BDQ resistance (Huitric et al., 2010). Therefore, new drugs that bind to unique sites on mycobacterial ATP synthase are needed. A recent screen led to discovery of a squaramide series of mycobacterial ATP synthase inhibitors (Tantry et al., 2017). The lead compound SQ31f (Fig. 1A, *orange*) inhibits mycobacterial ATP synthase with nanomolar IC_50_, high selectivity, and low cytotoxicity. This compound has shown promising ability to arrest the growth of *M. tuberculosis* in a mouse model of TB (Tantry et al., 2017) but how it binds the ATP synthase is unknown.

We determined electron cryomicroscopy (cryoEM) structures of *M. smegmatis* ATP synthase in complex with the second generation diarylquinoline TBAJ-876 and the squaramide lead compound SQ31f. TBAJ-876 binds to the same sites observed with BDQ in intact *M. smegmatis* ATP synthase (Guo et al., 2021), but with improved interaction of TBAJ-876’s tri- and dimethoxy pyridine rings with the enzyme’s a and c subunits compared to the aryl groups of BDQ. In contrast, SQ31f binds a unique site, buried at the interface between the c ring and the a subunit within the cytosolic half channel. Surprisingly, BDQ, TBAJ-876, and SQ31f all induce similar conformational changes in ATP synthase upon binding. Counter to previous claims (Sarathy et al., 2020), we show that at micromolar concentrations both TBAJ-587 and TBAJ-876, like BDQ, uncouple the PMF across the membrane of mycobacterial inner membrane vesicles. However, even micromolar BDQ does not restore ATP hydrolysis activity to ATP synthase when the enzyme is in a lipid bilayer, indicating that this observed behaviour with detergent-solubilized ATP synthase is unrelated to uncoupling of the PMF. High concentrations of SQ31f neither uncouple the PMF nor restore ATP hydrolysis activity to purified ATP synthase. SQ31f exhibits directional inhibition of the ATP synthase, with the IC_50_ for ATP hydrolysis approximately ten times higher than the IC_50_ for inhibition of ATP synthesis. Overall, this work reveals a new inhibitor binding site in mycobacterial ATP synthase and illuminates the nature of uncoupling, which may help guide development of compounds that kill mycobacteria by targeting the ATP synthase.

## Results

### Biochemical properties of diarylquinoline analogs

To investigate the activities of second generation diarylquinolines and squaramides, we obtained TBAJ-587-fumarate and TBAJ-876-tartrate as a gift from Dr. Christopher Cooper of TB Alliance (https://www.tballiance.org/) and synthesized the lead squaramide SQ31f following the reaction scheme outlined previously (Tantry et al., 2017). We then measured the ability of the compounds to inhibit ATP hydrolysis by detergent-solubilized and purified mycobacterial ATP synthase (Fig. 1B-*i*). ATP hydrolysis in wildtype mycobacterial ATP synthase is auto-inhibited by C-terminal extensions of the a subunits. These extensions prevent rotation of the rotor subcomplex in the direction necessary for ATP hydrolysis, which is opposite of its rotation direction during ATP synthesis (Guo et al., 2021). Therefore, the enzyme used in assays was activated by truncation of the a subunit extensions to allow ATP hydrolysis (Guo et al., 2021). The assays show that, like BDQ, TBAJ-587 and TBAJ-876 inhibit ATP hydrolysis at nanomolar concentrations but restore the hydrolytic activity of the enzyme at micromolar concentrations (Guo et al., 2021) (Fig. 1C, *blue, pink*, and *purple curves;* Fig. S1A). Loss of inhibition at high concentrations could occur if diarylquinolines disrupt the F_O_ region structure so that the inhibitor no longer blocks rotation of the rotor or ATP hydrolysis in the F_1_ region. If inhibitor-induced disruption of the F_O_ region also allowed protons to flow across the membrane independent of rotation and ATP synthesis, it would also explain why micromolar concentrations of BDQ have been reported to dissipate the PMF in mycobacteria and mycobacterial inverted membrane vesicles (IMVs) (Hards et al., 2018, 2015). The similar behavior of TBAJ-876 and BDQ was surprising because it was reported previously that TBAJ-876 does not induce uncoupling of the PMF (Sarathy et al., 2020) and we had assumed that micromolar uncoupling of the PMF and restoration of ATP hydrolysis activity were related.

In contrast to the diarylquinolines, SQ31f inhibits ATP hydrolysis with an IC_50_ of ~0.6 μM without restoring ATP hydrolysis at higher concentrations (Fig. 1C, *orange curve;* Fig. S1A). For the diarylquinolines, the extent to which ATP hydrolysis activity is restored appears to correlate with the hydrophobicity of the compounds, with the most hydrophobic drug, BDQ, restoring the most ATP hydrolysis activity (Fig. 1C, *blue curve*) (Guo et al., 2021), and the most hydrophilic compound, TBAJ-876, restoring the least activity (Fig. 1C, *purple curve*). To determine if this restoration of ATP hydrolysis activity also occurs when ATP synthase is in a lipid bilayer, we measured ATP hydrolysis by inverted membrane vesicles (IMVs) from the *M. smegmatis* strain with truncated a subunits (Fig. 1B-*ii*). In this assay, high concentrations of even the most hydrophobic inhibitor, BDQ, did not restore ATP hydrolysis activity (Fig. 1D, Fig. S1B). Together, these results indicate that restoration of ATP hydrolysis by detergent-solubilized ATP synthase at high concentrations of diarylquinolines is an artefact that occurs when the enzyme is in a detergent micelle rather than a lipid bilayer and is not related to uncoupling of the PMF in mycobacteria and mycobacterial IMVs. We next evaluated the ability of BDQ, TBAJ-587, and TBAJ-876 to uncouple a transmembrane PMF built up across the membranes of IMVs (Fig. 1B-*iii*). In this assay (Hards et al., 2015), IMVs are incubated with the fluorophore 9-amino-6-chloro-2-methoxyacridine (ACMA) or *N,N,N’,N’*-tetrametlivlacridine-3,6-diamine (acridine orange), depending on the absorption maxima of the putative uncoupling compound being tested. Addition of succinate allows succinate dehydrogenase from the IMVs to reduce endogenous menaquinone, which is then oxidized by the respiratory CIII2CIV2 supercomplex and cytochrome *bd*. Both CIII2CIV2 and cytochrome *bd* translocate protons into the IMVs, acidifying the IMVs, which traps and concentrates the fluorophore and quenches its fluorescence. Addition of the ionophore nigericin to acidified IMVs results in dissipation of the PMF (Fig. 1E, *cyan curve*). As described previously, addition of 20 μM BDQ also dissipates the PMF (Hards et al., 2015) (Fig. 1E, *blue curve*), although more slowly than nigericin. Counter to previous reports (Sarathy et al., 2020), addition of either TBAJ-876 or TBAJ-587 also dissipates the PMF (Fig. 1E, *pink* and *blue curves*), although more slowly than either BDQ or nigericin. The uncoupling activities of BDQ, TBAJ-587, and TBAJ-876 occur in a dose dependent manner (Fig. S1C). These experiments show that in addition to ATP synthase inhibition, TBAJ-587 and TBAJ-876 retain the uncoupling activity of BDQ. In *M. tuberculosis* BDQ activates the mycobacterial dormancy regulon, inducing metabolic changes including a reliance on glycolysis that avoids bacteriolysis for several days (Koul et al., 2014). The complex mechanism of killing by BDQ has been proposed to rely on both its inhibition of ATP synthesis and its uncoupling of the PMF (Hards et al., 2018, 2015).

### TBAJ-876 binds ATP synthase with improved interactions compared to BDQ

To determine if the second generation diarylquinoline TBAJ-876 binds ATP synthase in the same way as BDQ, we incubated the enzyme with 200 μM TBAJ-876, prepared cryoEM specimens, and subjected the sample to structure determination by cryoEM (Fig. 2 and Fig. S3). This analysis resulted in three-dimensional (3D) maps of the three rotational states of the enzymes at resolutions ranging from 2.6 to 2.9 Å (Fig. S3, Table 1). Focused refinement of the F_O_ region of the complex resulted in a map at 2.8 Å resolution (Fig. S3, Table 1). Together, these maps allowed construction of atomic models for intact ATP synthase in three rotational states with TBAJ-876 bound (Table 1).

**Figure 2.**
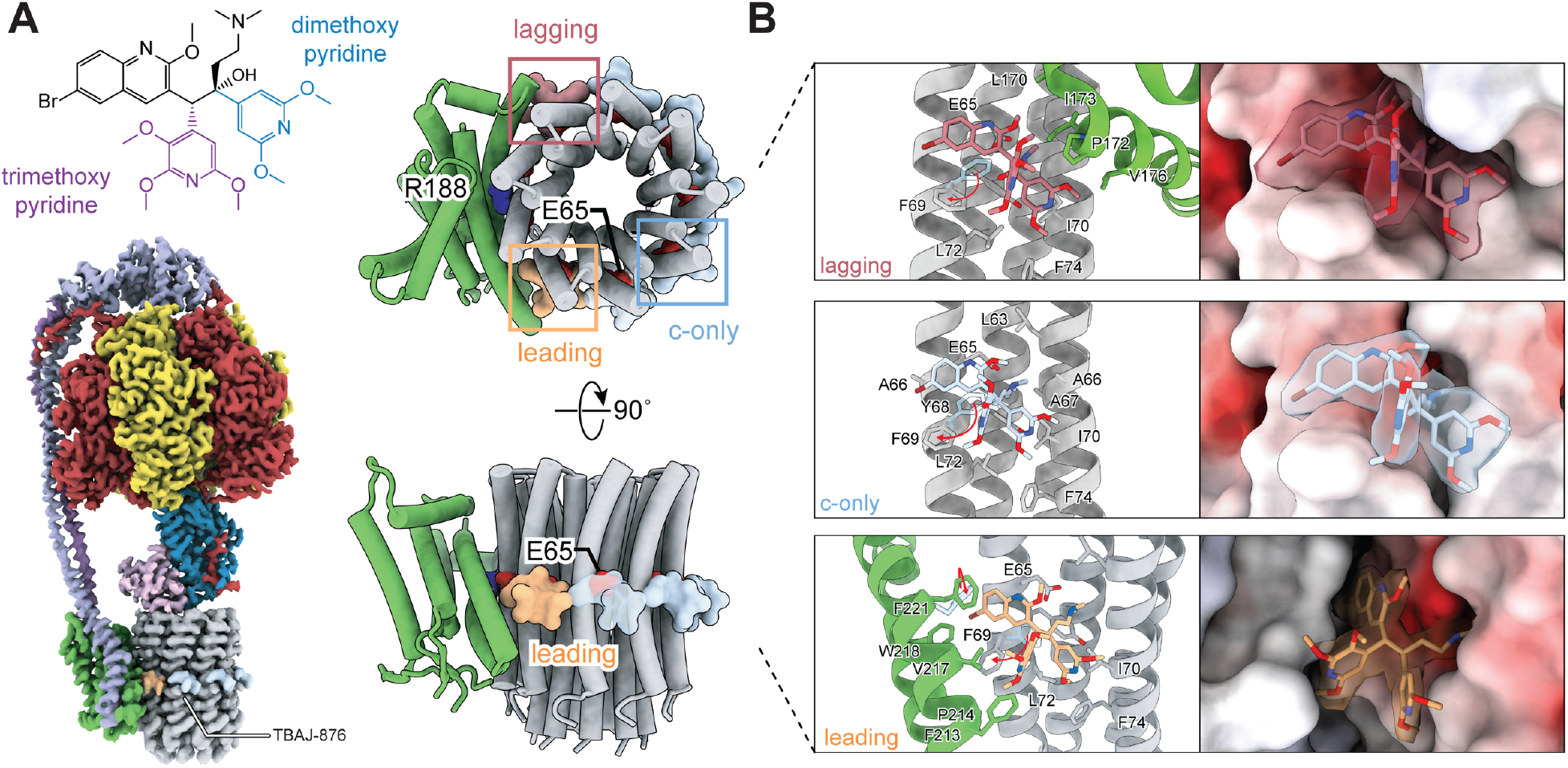
Structure of *M. smegmatis* ATP synthase bound to TBAJ-876. **A,** TBAJ-876 binds five c-only sites (*blue*), a leading site (*orange*), and a lagging site (*red*). **B,** Atomic model (*left*) and surface representation (*right*) of TBAJ-876 in the lagging site (*top*), c-only site (*middle*), and leading site (*bottom*). Red arrows indicate movement of residues upon binding of TBAJ-876, with their drug-free positions indicated in semi-transparent light blue. Residues that form close interactions with the inhibitor are indicated.

The resolution of the cryoEM maps with TBAJ-876 compare favorably with previous maps of ATP synthase with BDQ (Guo et al., 2021; Montgomery et al., 2021), and may be valuable for medicinal chemistry efforts. As seen with saturating concentrations of BDQ (Guo et al., 2021), the maps show seven TBAJ-876 molecules bound to the F_O_ region (Fig. 2A). Five of the TBAJ-876 molecules bind at the “c-only” sites that involve just subunit c (Fig. 2A, *light blue surfaces*). The two additional TBAJ-876 molecules bind at the “leading” (Fig. 2A, *orange surface*) and “lagging” (Fig. 2A, *red surface*) sites at the interface of subunits a and c. Many aspects of the interaction of TBAJ-876 with the enzyme are conserved with BDQ (Guo et al., 2021). Briefly (Fig. 2B, *left*), in all three sites, Phe69 undergoes a conformational change from the drug-free structure to provide a hydrophobic platform, allowing the inhibitor to insert between two c subunits. TBAJ-876’s dimethylamino group forms a salt bridge with the carboxyl group of the proton carrying Glu65. Tyr68 and Ala66 from one c subunit and Leu63, Ala66, Ala67, and Ile70 from another c subunit form Van der Waals contacts with TBAJ-876. Compared to BDQ, the extended π-system from TBAJ-876’s trimethoxy pyridine ring provides improved interactions with Phe69 (Fig. S4, *middle*). Similarly, the dimethoxy pyridine ring provides improved contact with Ile70, Leu72, and Phe74 in the c ring (Fig. S4, *middle*). As with BDQ, the lagging site Leu170, Pro172, Ile173, and Val176 from subunit a interact with TBAJ-876, with the trimethoxy pyridine forming improved contacts with Pro172 compared to BDQ (Fig. 2B, *top*, Fig. S4, *top*). Similar to BDQ, the leading site Phe213, Pro214, Val217, Trp218, and Phe221 from subunit a interact with TBAJ-876, and Phe221 undergoes a conformational change to accommodate insertion of the diarylquinoline moiety at the a-c interface (Fig. 2B, *bottom*). However, in the leading site, the trimethoxy pyridine ring provides improved interactions with Phe213, Pro214, and Val217 (Ala211, Pro212, and Ile215 in *M. tuberculosis*, respectively) of the a subunit (Fig. S4, *bottom*). These increased interactions with TBAJ-876 compared to BDQ could help explain the 10-fold improvement in MIC for TBAJ-876 (Sutherland et al., 2019) and improved IC_50_ in ATP synthesis assays with *Mycobacterium bovis* IMVs (Sarathy et al., 2019). The ATP hydrolysis assays performed here used 2 nM enzyme, which prevents measurement of IC_50_ values below 1 nM, and consequently could not detect a decreased IC_50_ of TBAJ-876 compared to BDQ. As with previous structures of mycobacterial ATP synthase with BDQ (Guo et al., 2021; Montgomery et al., 2021) we found no evidence of diarylquinolines binding to the ε subunit, which was suggested previously (Kundu et al., 2016; Sarathy et al., 2019).

### Squaramide inhibits ATP synthesis more potently than ATP hydrolysis

To investigate how inhibition with squaramides differs from inhibition with diarylquinoline, we performed assays with SQ31f. Using *M. smegmatis* grown in liquid culture, we measured a MIC_90_ for SQ31f of 12.5 μM, which is higher than the MIC80 of 0.5 μM reported previously for *M. tuberculosis* (Tantry et al., 2017). Neither SQ31f nor BDQ had bactericidal activity against *M. smegmatis* at up to 8× the MIC_90_ (Fig. S5). When added to IMVs acidified by proton pumping by the electron transport chain (Fig. 1B-*iii*), SQ31f did not uncouple the PMF at concentrations up to 80 μM (Fig. 3A, Fig. S1D), consistent with previous reports (Tantry et al., 2017). As described above, the IC_50_ for inhibition of ATP hydrolysis by SQ31f was ~0.6 μM (Fig. 3B, *blue curve*). This value was much higher than the 30 nM IC_50_ for inhibition of ATP synthesis by *M. smegmatis* IMVs reported previously (Tantry et al., 2017). Therefore, we performed similar ATP synthesis assays where we established a PMF in wildtype *M. smegmatis* IMVs and monitored ATP synthase activity with luciferase (Lemasters and Hackenbrock, 1978; Preiss et al., 2015) (Fig. 1B-*iv*). In these assays, the IC_50_ for SQ31f inhibition of ATP synthesis was 38 nM (Fig. 3B, *purple curve*), similar to previous results (Tantry et al., 2017). Therefore, the IC_50_ for inhibition of ATP synthesis by SQ31f appears to be more than ten times lower than the IC_50_ for inhibition of ATP hydrolysis. While this discrepancy may be due in part to experimental differences in the different assays, it suggests that SQ31f is a somewhat directional inhibitor, blocking rotation of the rotor subcomplex in the ATP synthesis direction more effectively than it blocks rotation of the rotor in the ATP hydrolysis direction. Asymmetric inhibition highlights the importance of performing ATP synthesis assays when detecting ATP synthase inhibitors in high-throughput screens.

**Figure 3.**
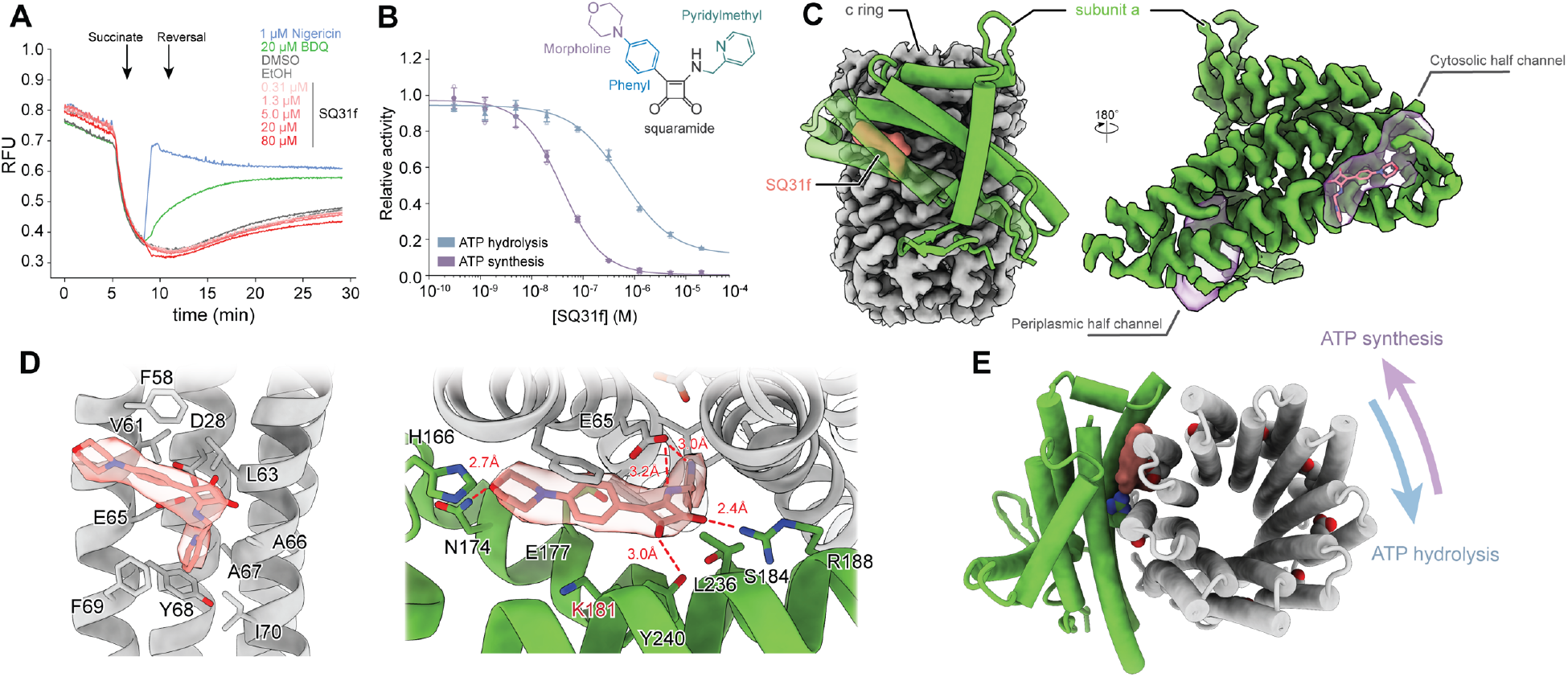
Inhibition of *M. smegmatis* ATP synthase by SQ31f. **A,** Uncoupling activity of SQ31f in *M. smegmatis* IMVs monitored with acridine orange fluorescence. **B,** SQ31f inhibition of ATP hydrolysis by purified ATP synthase with truncated α subunits (blue curve) occurs at higher concentrations than inhibition of ATP synthesis by *M. smegmatis* IMVs (purple curve). Mean relative activity (filled symbols) ± s.d from n=3 separate assays (triangle and circles). **C,** SQ31f (red surface and model) binds at the subunit a-c interface in the cytosolic proton half channel. Half channel cavities (purple surfaces) were calculated with *Hollow* (Ho and Gruswitz, 2008). **D,** Interaction between SQ31f and subunits a and c in the F_O_ region. Putative hydrogen bonds are shown as red dashed lines, with distances between participating atoms indicated. **E,** Atomic model of SQ31f bound to the F_O_ region of ATP synthase. The direction of c ring rotation is indicated for ATP hydrolysis (blue arrow) and ATP synthesis (purple arrow).

### Squaramide-binding reveals a new inhibitor pocket on ATP synthase

To understand the structural basis for the unusual inhibition of ATP synthase by SQ31f, we incubated the enzyme with the inhibitor and prepared specimens for cryoEM. This analysis provided maps of the three rotational states of the enzymes at nominal overall resolutions of 2.8 to 3.1 Å, with focused refinement of the F_O_ region resulting in a map at 2.8 Å resolution (Fig. S2, Fig. S3, Table 1). Together, these maps allowed construction of atomic models for intact ATP synthase in three rotational states with SQ31f bound (Table 1). The cryoEM maps show that SQ31f binds a single site in the F_O_ region of the complex (Fig. 3C, *left*). The squaramide binding site is buried at the subunit a-c interface, consistent with resistance to SQ31f in *M. tuberculosis* where Lys179 in subunit a (equivalent to Lys181 in *M. smegmatis*, Fig. 3D, *right*) is mutated to asparagine (Tantry et al., 2017). SQ31f fits tightly in the cavity corresponding to the cytosolic proton half channel, where protons normally exit the enzyme during ATP synthesis (Fig. 3C, *right*). The cytosolic half channel is expected to be filled with water (Guo et al., 2017, 2019), consistent with the high aqueous solubility of SQ31f (cLogP=1.02) compared to TBAJ-876 (cLogP=5.15) and other diarylquinolines. As a result of binding in this aqueous cavity, interaction of SQ31f with the enzyme is mediated by numerous polar residues. The pyridine and secondary amine group of SQ31f appear to form hydrogens bonds with Glu65, as predicted by structure-activity relationship studies (Tantry et al., 2017), with the aromatic pyridine ring forming additional contacts with Phe69 and Tyr68 from one c subunit and Ile70, Ala66, and Ala67 from another c subunit (Fig. 3D, *left*). The carbonyl groups from the squaramide scaffold form apparent hydrogen bonds with Arg188 and Tyr240 from subunit a, and additional Van Der Waals and polar interactions with Leu63 from subunit c, as well as Leu236 and Ser184 from subunit a (Fig. 3D, *right*). The phenyl ring of SQ31f interacts with Val61 and Phe58 from a c subunit, as well as Glu177 and Lys181 from the a subunit. Lastly, the SQ31f morpholine ring interacts with Asn174 and His166 from subunit a, with Asn174 forming an apparent hydrogen bond with either the protein backbone carbonyls from subunit a or the morpholine ether in SQ31f (Fig. 3D, *right*). The observed binding site for SQ31f may explain its directional inhibition: rotation of the rotor in the hydrolysis direction pulls the compound away from the a subunit (Fig. 3E, *blue arrow*), which would break many of the interactions that hold it in place. In this way, ATP hydrolysis could help dislodge the inhibitor from the cytosolic half channel. In contrast, rotation of the rotor in the ATP synthesis direction (Fig. 3E, *purple arrow*) pushes the compound deeper into its binding site, blocking rotation and inhibiting ATP synthesis.

### Diarylquinoline and squaramide binding induce similar conformational changes

BDQ, TBAJ-876, and SQ31f all induce large conformational changes in the mycobacterial ATP synthase compared to the inhibitor free structure (Fig. 4A, Video 1). On inhibitor binding, minimal movement occurs between the catalytic α3β3 hexamer and the γεc9 rotor. However, binding of any of the three inhibitors causes the rotor subcomplex and α3β3 hexamer to rotate together relative to subunits a, b, and b-δ. This rotation twists the δ region of the b-δ subunit. Remarkably, the amount of rotation is similar for all three inhibitors. Comparable to BDQ, TBAJ-876 binding causes a ~28° rotation in the synthesis direction for rotational State 1, a ~30° rotation in the synthesis direction for State 2, and a ~12° rotation in the hydrolysis direction for State 3. Similarly, SQ31f binding leads to a ~24° rotation in the synthesis direction for State 1, a ~25° rotation in the synthesis direction in State 2, and a ~16° rotation in the hydrolysis direction for State 3 (Fig. 4B). For the diarylquinolines, these rotations are necessary for simultaneous creation of the high affinity leading and lagging sites, whereas for squaramide the rotations align Glu65 with the cytosolic half channel opening to allow inhibitor binding.

**Figure 4.**
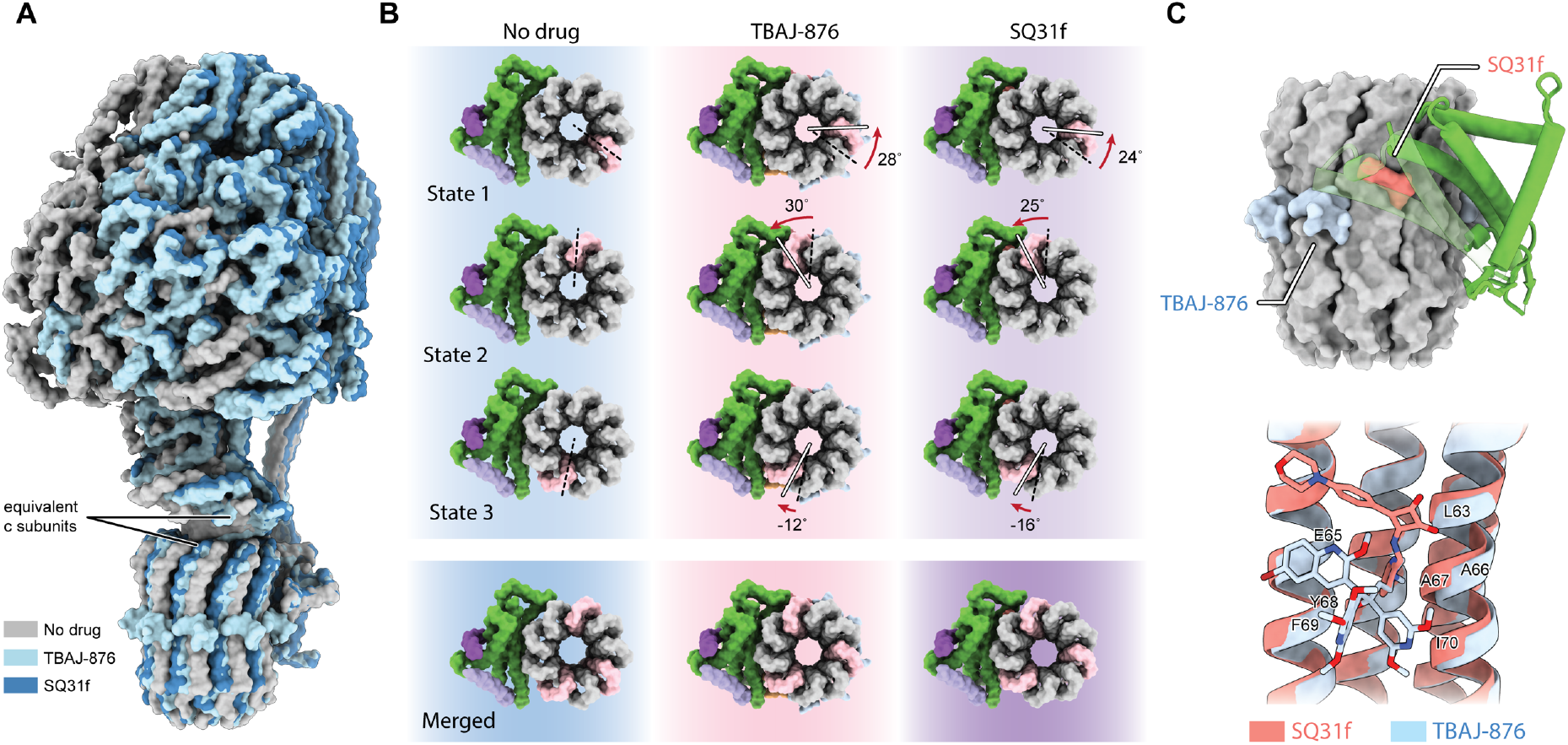
Diarylquinoline and squaramide binding both induce a symmetric distribution of rotational states. **A,** TBAJ-876 and SQ31f induce rotation of the α3β3 hexamer and c9εγ rotor relative to subunits a, b, and b-δ. **B,** The magnitude and direction of rotation in each of the three states in the presence of TBAJ-876 or SQ31f is indicated, with positive rotations used for the synthesis direction and negative rotations used for the hydrolysis direction. Equivalent c subunits (pink) are defined by their position relative to the ε subunit. Overlay of the c ring position from each rotational state shows that inhibitor binding causes the states to become three-fold symmetric. **C,** Comparison of the SQ31f and TBAJ-876 binding sites (top), and interactions with the c ring (bottom).

The *M. smegmatis* ATP synthase possesses a nine-fold symmetric c ring that matches the pseudo-three-fold symmetry of the α3β3 hexamer in the F1 region. Therefore, one might expect the rotational states of the inhibitor-free structures to differ by 120° steps in the F_O_ region, just as these states differ by 120° steps in the F_1_ region. However, as shown previously (Guo et al., 2021; Montgomery et al., 2021), in the inhibitor-free structures, rotational states 1, 2, and 3 are separated by 120°, 160°, and 80° steps in the *F*_O_ region (Fig. 2B, *left*). In the inhibitor-bound structures, the rotation in the synthesis direction for States 1 and 2 are slightly smaller with SQ31f than with TBAJ-876, while the rotation in the hydrolysis direction for State 3 is slightly larger with SQ31f than with TBAJ-876. These rotations lead the F_O_ region conformations to differ by ~120° rotations between states for both types of inhibitors, restoring symmetry to the complex. Whether or not these symmetric conformations are due to local energy minima that favor inhibitor binding is unclear. Nonetheless, the similarity in conformation of the inhibited states with both classes of inhibitor suggests that a computational search for binding pockets for new inhibitors should consider the conformations seen with diarylquinolines and squaramides.

## Discussion

SQ31f and diarylquinolines both inhibit the mycobacterial ATP synthase by preventing rotation of the c ring relative to subunit a in the F_O_ region of the enzyme. However, the structures reported here show that squaramides and diarylquinolines bind distinct sites, with only some of the same residues mediating interaction with the c ring (Fig. 4C). The diarylquinolines BDQ, TBAJ-587, TBAJ-876 bind in mostly hydrophobic pockets. Conversely, SQ31f binds in the aqueous environment of the cytosolic half channel. This difference in binding site allows SQ31f to retain its efficacy against BDQ-resistant *M. tuberculosis* strains harboring A63P or I66M mutations in their c subunits (Tantry et al., 2017). Cross-resistance may nonetheless appear in some *M. tuberculosis* strains harboring D28N mutations (D32 in *M. smegmatis*) in the c subunit (Tantry et al., 2017). The D28N mutation disrupts the hydrogen-bonding network that stabilizes the Glu65 conformation needed for binding of diarylquinolines (Preiss et al., 2015) and also disrupts squaramide binding (Tantry et al., 2017), presumably by the same mechanism.

Although increasing the number of hydrophilic interactions of SQ31f with the mycobacterial ATP synthase should enhance SQ31f binding affinity, the addition of polar moieties may limit the compound’s ability to cross the mycobacterial inner membrane to reach its target. How the hydrophilic character and overall structure of SQ31f affect hERG channel inhibition, sensitivity to mmpl5/mmps5 drug efflux, and overall toxicity compared to diarylquinolines remains to be investigated.

As shown above, BDQ, TBAJ-587, and TBAJ-876 can all transport ions across the lipid bilayer in mycobacterial inner membrane vesicles to uncouple the PMF. The uncoupling property of BDQ has been observed *in vitro* at micromolar concentrations and was proposed to be important for the drug’s ability to kill mycobacteria *in vivo* (Andries et al., 2005; Hards et al., 2015). In contrast, SQ31f lacks detectable uncoupling activity *in vitro* (Fig. 3A) and shows only a bacteriostatic effect on *M. tuberculosis in vivo* (Tantry et al., 2017). It is not clear whether *in vitro* uncoupling activity is necessary for bacterial killing in mouse models of *M. tuberculosis* infection. However, if it is, the screen that led to the discovery of SQ31f, which excluded compounds with pronounced uncoupling activities due to concerns about toxicity, could have excluded some potentially valuable compounds (Tantry et al., 2017). Therefore, elucidating the mechanism of uncoupling, determining whether it synergizes with ATP synthase inhibition for killing of *M. tuberculosis*, and establishing whether it is beneficial *in vivo*, could provide important insight needed for the discovery of new antimycobacterial compounds.

## Methods

### Synthesis of SQ31f

3-(4-Morpholinophenyl)-4-((pyridin-2-ylmethyl) amino)-cyclobut-3-ene-1,2-dione (SQ31f) was synthesised as described previously (Tantry et al., 2017). A mixture of 3,4-dihydroxycyclobut-3-ene-1,2-dione (5 g, 43.84 mmol) and thionyl chloride (6.7 mL, 92.06 mmol) was cooled to 0 °C. Under vigorous stirring, dimethylformamide (15 drops) was added and the mixture was refluxed at 80 °C for 3 h. The reaction mixture was cooled to room temperature, and excess thionyl chloride was evaporated under reduced pressure. 3,4-Dichlorocyclobut-3-ene-1,2-dione was obtained as a residual yellow solid and used without further purification. The 3,4-dichlorocyclobut-3-ene-1,2-dione (2.00 g, 13.25 mmol) was dissolved in dried dichloromethane (30 ml) and cooled to 0 °C. Anhydrous AlCl_3_ (1.77 g, 13.25 mmol) was added carefully. N-phenylmorpholine (1.43 g, 8.77 mmol) was added in portions to the reaction. After completion of addition, the mixture was warmed to room temperature and stirring was continued for additional 2 h. The mixture was quenched with cold water and the crude product was extracted with dichloromethane (2 × 30 mL). The combined organic layer was washed with water (20 mL), brine (20 mL), dried over Na_2_SO_4_, and evaporated under reduced pressure. After purification by medium pressure liquid chromatography (MPLC) with silica gel column (ethyl acetate/hexane gradient), 3-chloro-4-(4-morpholinophenyl)-cyclobut-3-ene-1,2-dione was obtained as a red solid (17 % yield, 2.28 mmol, 0.63 g). The 3-chloro-4-(4-morpholinophenyl)-cyclobut-3-ene-1,2-dione (0.30 g, 1.08 mmol) was dissolved in dioxane (30 mL). Triethylamine (0.225 mL) was added, followed by addition of 2-picolylamine (0.167 mL, 1.62 mmol). The reaction was stirred for 2 h and solvent was evaporated under reduced pressure. The residue was dissolved in ethyl acetate and washed with water (20 mL) and brine (20 mL) and dried over Na2SO4. After evaporating the solvent and purifying with MPLC using a silica gel column (methanol/dichloromethane gradient), 0.25 g of 3-(4-morpholinophenyl)-4-((pyridin-2-ylmethyl)-amino)-cyclobut-3-ene-1,2-dione) was obtained as a yellow solid (66 % yield, 0.72 mmol).

**^1^H-NMR** (400 MHz, DMSO-d_6_): δ 9.36 (t, *J* = 6.3 Hz, 1H), 8.53 (ddd, *J* = 4.9, 1.8, 0.9 Hz, 1H), 8.02 – 7.89 (m, 2H), 7.79 (td, *J* = 7.7, 1.8 Hz, 1H), 7.41 (dt, *J* = 7.8, 1.0 Hz, 1H), 7.30 (ddd, *J* = 7.6, 4.8, 1.1 Hz, 1H), 7.10 – 6.96 (m, 2H), 4.99 (d, *J* = 6.2 Hz, 2H), 3.73 (t, *J* = 4.9 Hz, 4H), 3.27 (t, *J* = 4.9 Hz, 4H) ppm;

**^13^C-NMR** (100 MHz, DMSO-dő): δ 192.22, 189.43, 178.51, 163.32, 157.85, 152.75, 149.68, 137.48, 128.41, 123.10, 122.01, 119.77, 114.34, 66.30, 49.25, 47.38 ppm; **MS** (APCI, MeOH): m/z 350.1 ([M+H]^+^), calc: 349.14; **HPLC purity**: 99.0%.

### Bacterial growth and protein purification

*Mycobacterium smegmatis* strain GMC_MSM1, which includes a 3×FLAG tag at the C terminus of the β subunits, and GMC_MSM2, with a 3×FLAG tag truncating the a subunits at residue S518 (Guo et al., 2021), were grown in Middlebrook 7H9 medium supplemented with 0.8 g/L NaCl, 2 g/L dextrose, 10 g/L tryptone, and 0.5 mL/L Tween 80. Cells from 6 L cultures grown for 48 h were collected by centrifugation (6,500 g, 20 min) and resuspended in 150 mL lysis buffer (50 mM Tris-HCl pH 7.5, 150 mM NaCl, 5 mM MgSO4, 5 mM 6-aminocaproic acid, 5 mM benzamidine hydrochloride, and 1 mM PMSF). Resuspended cells were filtered with Miracloth (Millipore) and lysed using an Emulsiflex-C3 High Pressure Homogenizer (Avestin) with three passages at 20 kpsi. Unbroken cells and debris were removed by centrifugation at 39,000 g for 30 min. To collect membranes, the supernatant was centrifugated at 200,000 g for 60 min, and membranes were resuspended in membrane buffer (50 mM Tris-HCl pH 7.4, 150 mM NaCl, 15% [v/v] glycerol, 5 mM MgSO4, 5 mM 6-aminocaproic acid, 5 mM benzamidine hydrochloride, and 1 mM PMSF), flash frozen in liquid nitrogen, and stored at −80°C.

Thawed membranes were solubilized with 1% (w/v) dodecyl-B-D-maltoside (DDM) (Anatrace). Insoluble material was removed by centrifugation at 200,000 g for 60 min. Solubilized membranes were filtered with a 0.45 μM filter and loaded on a 2 mL column of M2 affinity matrix (Sigma) equilibrated with wash buffer (50 mM Tris-HCl pH 7.4, 150 mM NaCl, 15% [v/v] glycerol, 5 mM 6-aminocaproic acid, 5 mM benzamidine hydrochloride, and 0.05% [w/v] DDM). After washing with ~10 column volumes of wash buffer, bound proteins were eluted with three column volumes of wash buffer with 150 μg/mL 3×FLAG peptide. The sample was concentrated with a 100 kDa cutoff concentrator to 500 μL and loaded on a Superose 6 Increase 10/300 column (GE Healthcare) equilibrated with gel filtration buffer (20 mM Tris-HCl pH 7.4, 100 mM NaCl, 15% [v/v] glycerol, and 0.05% [w/v] DDM). Eluted ATP synthase was concentrated to ~6 mg/mL and used for cryoEM sample preparation, or flash frozen and stored at −80°C for assays.

### Biochemical assays

NADH coupled ATP hydrolysis assays with detergent solubilized ATP synthase were performed as described previously (Guo et al., 2021). Briefly, 2 nM hydrolytically competent ATP synthase (from strain GMC_MSM2) was assayed in 160 μL ATPase buffer (50 mM Tris-HCl pH 7.4, 150 mM NaCl, 10% [v/v] glycerol, 5 mM MgCl_2_, 0.2 mM NADH, 3.2 units pyruvate dehydrogenase, 8 units lactate dehydrogenase, and 0.05% [w/v] DDM). The reaction was initiated by addition of ATP to 2 mM and 1 mM phopshoenol pyruvate and the change in absorbance at 340 nm over time was monitored with a Neo2 Plate reader (Agilent). Assays were performed at 25°C.

For ATP hydrolysis assays with IMVs, endpoint measurements of phosphate concentration were performed with malachite green (Hess and Derr, 1975; Lanzetta et al., 1979; Baykov et al., 1988). The malachite green dye solution contains 0.44 g malachite green in 3 M sulfuric acid. Before use, 2.5 mL of 7.5% ammonium molybdate and 0.2 mL of 11% tween 20 were added to 10 mL of the dye solution to obtain the colour reagent (Baykov et al., 1988). GMC_MSM2 *M. smegmatis* cells were lysed in malachite lysis buffer (50 mM Tris-HCl pH 7.5, 5 mM MgCl_2_, 5 mM 6-aminocaproic acid, 5 mM benzamidine hydrochloride, and 1 mM PMSF). Unbroken cells and debris were removed by centrifugation at 39,000 g for 30 min. Membranes were collected by centrifugation at 200,000 g for 60 min and resuspended in malachite IMV buffer (50 mM Tris-HCl pH 7.5, 5 mM MgCl_2_, 10% [v/v] glycerol, 5 mM 6-aminocaproic acid, 5 mM benzamidine hydrochloride) to form IMVs. IMVs were diluted to 80 μL at 0.2 mg/mL total protein concentration with IMV ATPase buffer (10 mM HEPES-KOH pH 7.5, 100 mM KCl, 5 mM MgCl_2_) and the reaction started with 2 mM ATP. After 40 min, 20 μL from the reactions was diluted by adding 140 μL MilliQ water and the reaction stopped by flash freezing. To assay inorganic phosphate production, samples were thawed and 40 μL colour reagent (Baykov et al., 1988) was added to the diluted samples. After 10 min, the colour reaction was stopped by adding 23 μL of 25% (w/v) sodium citrate. Finally, absorbance was read at 630 nm with a Spectramax M5e plate reader (Molecular Devices). Assays were performed at room temperature. ATP hydrolysis activity by the ATP synthase was defined as hydrolysis activity that was sensitive to 100 μM DCCD.

ATP synthesis assays were performed with membranes harvested from GMC_MSM1 as described previously (Lemasters and Hackenbrock, 1978; Preiss et al., 2015; Hards et al., 2018). Briefly, GMC_MSM1 *M. smegmatis* cells were lysed in lysis buffer and membranes resuspended in membrane buffer to form IMVs. IMVs were diluted to 160 μL at 0.5 mg/mL total protein concentration in ATP synthesis buffer (20 mM HEPES pH 7.0, 100 mM potassium acetate, 10 mM Na2HPO4, 5 mM magnesium acetate, 80 μg/mL luciferase, 400 μM D-luciferin, 50 μM ADP, 200 μM P1,P5-Di[adenosine-5’]pentaphosphate [Ap5A]) and incubated for 1 h at room temperature. The reaction was started by addition of 5 mM sodium succinate (pH ~8) or water (control) and luminescence monitored with a Neo2 plate reader with a luminescence filter cube. ATP was added to 10 μM from a 117 μM stock in each well at the end of each run to serve as an internal standard. The ATP concentration at the completion of each reaction was calculated using the internal standards to determine relative activities (Lemasters and Hackenbrock, 1978). Assays were performed at 25°C.

Proton pumping assays (Hards et al., 2015) were performed with IMVs harvested from GMC_MSM1, as described above for ATP synthesis assays. Briefly, IMVs were diluted to 160 μL at 0.5 mg/mL total protein concentration in IMV ACMA buffer (10 mM HEPES-KOH pH 7.5, 100 mM KCl, 5 mM MgCl_2_, 3 μM ACMA). For assay of SQ31f, IMV Acridine orange buffer was used instead of IMV ACMA buffer because of overlapping absorbance peaks for ACMA and SQ31f (10 mM HEPES-KOH pH 7.5, 100 mM KCl, 5 mM MgCl_2_, 5 μM acridine orange). Reactions were started with 5 mM sodium succinate (~pH 8). Inhibitors were added after full acidification of the vesicles to measure their protonophoric activity. Assays were performed at 37°C. For all assays, inhibitors were diluted to a final DMSO concentration of 2% in the reaction buffer. Fluorescence was excited at 410 nm and monitored at 480 nm for ACMA or excited at 493 nm and monitored at 530 nm for acridine orange.

### MIC determination

MICs were determined against *M. smegmatis* mc^2^ 155 pTEC27 (Takaki et al., 2013) (a gift from Dr. Lalita Ramakrishnan: Addgene plasmid #30182; http://n2t.net/addgene:30182; RRID: Addgene_30182) by the broth microdilution method as described previously (Richter *et al*. 2018). Briefly, 96-well flat bottom tissue culture plates (Sarstedt, 83.3924.500) were used. In the third well of each row, two times the highest desired concentration of the inhibitor was added in 7H9 medium supplemented with 10% ADS and 0.05% polysorbate 80. Each compound was diluted twofold in a nine-point serial dilution. The concentration of the starting inoculum was 5 × 10^5^ cells/mL. The starting inoculum was diluted from a preculture at the mid-log phase (OD_600_ 0.3 to 0.7) and an OD_600_ of 0.1 was correlated to 1 × 10^8^ colony forming units (CFU/mL). The plates were sealed with parafilm, placed in a container with moist tissue, and incubated for three days at 37°C. Each plate had eight negative controls (1% DMSO) and eight positive controls (100 μM amikacin). After incubation the plates were monitored by OD measurement at 590 nm (BMG Labtech Fluostar Optima). The assay was performed in duplicate and results were validated by measurement of red fluorescent protein. Every assay plate contained eight wells with 1% DMSO as negative controls, which corresponds to 0% growth inhibition, and eight wells with 100 μM amikacin as positive controls, which corresponds to 100% growth inhibition. The lowest concentration of sample with > 90% growth inhibition is reported as the MIC_90_. Growth inhibition was calculated with:

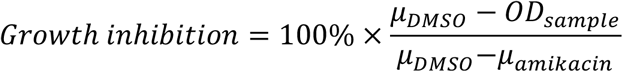

where *μ_DMSO_* and *μ_amikacin_* are the mean OD for wells with DMSO and amikacin, respectively, and *OD_sample_* is the OD of the well with sample compound. Assay quality was monitored through determination of the *Z*’ score, with:

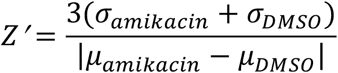

where σ_amikacin_ and σ_SDMSO_ are the standard deviation of the OD for wells with amikacin and DMSO, respectively. The *Z*’ score for each assay was above 0.4.

### Minimum bactericidal concentration determination

For minimum bactericidal concentration (MBC) determination against *M. smegmatis mc^2^ 155 pTEC27* was incubated in a microplate dilution assay as described above for MIC determination.

Subsequently, the MBC was determined by CFU counting. The wells of 6 well plates were filled with 4 mL 7H10 agar supplemented with 0.5% glycerol, 10% ADS, and 400 μg/mL hygromycin. From the drug concentrations where growth inhibition was detected in the microplate dilution assay, 10 μL (undiluted or diluted 1:20 or 1:100) were plated onto wells of the 6 well plates. Colonies were counted after three days of incubation at 37°C and the experiment was carried out in triplicate with CFUs/mL calculated from the results. The number of CFUs was also determined in the inoculum prior to the three-day incubation.

### CryoEM

Homemade holey gold grids were nanofabricated as previously described (Marr et al., 2014). For structure determination with TBAJ-876, 200 μM TBAJ-876 was added to purified ATP synthase at 6 mg/mL and incubated for 5 min, before removing glycerol with a Zeba spin desalting column (Thermo Scientific) equilibrated with cryoEM buffer (20 mM Tris-HCl pH7.4, 100 mM NaCl, and 0.05% [w/v] DDM) containing 200 μM TBAJ-876 (2% DMSO). For the SQ31f specimen, glycerol was removed from the ATP synthase sample first, and the enzyme was subsequently incubated with 100 μM inhibitor for 5 min. Grids were glow discharged in air for 2 min and sample (2 μL) applied in an EM GP2 plunge freezer (Leica) at 4°C and 90% humidity. Grids were blotted for 1 s before freezing in liquid ethane.

### Data collection

CryoEM movies were collected on a 300 kV Titan Krios G3 electron microscope with a Falcon 4i camera (Thermo Fisher Scientific). Automated data collection was performed with EPU. For both datasets, movies were collected at 75,000× magnification, with a calibrated pixel size of 1.03 Å. For the TBAJ-876 dataset, 8,112 movies were collected with an exposure rate of 7 e^-^ /pixel/s and a total exposure of ~44 e^-^/Å^2^. For the SQ31f dataset, 4,012 movies were collected with an exposure rate of 6.9 e^-^/pixel/s and a total exposure of ~54 e^-^/Å^2^.

### Image analysis

Except where specified, cryoSPARC v.3 was used for image analysis (Punjani et al., 2017) and the TBAJ-876 and SQ31f datasets were processed the same way (Fig. S2). Exposure fractions were aligned with MotionCor2 (Zheng et al., 2017). Template picking yielded 1,281,206 particle images for the TBAJ-876 dataset and 805,534 particle images for SQ31f dataset. Datasets were cleaned with two rounds of 2D classification, yielding 639,028 particle images for TBAJ-876 and 288,126 for SQ31f. Ab initio 3D classification and heterogenous refinement yielded the three rotational states of the ATP synthase, which were refined further with nonuniform refinement (Punjani et al., 2020) and CTF refinement. Particle image parameters were converted to Relion 3.0 .star format with *pyem* to correct for beam-induced motion using Bayesian polishing (Zivanov et al., 2019) before being reimported to cryoSPARC. Further CTF refinement and non-uniform refinement yielded the final maps of ATP synthase in its three main rotational states. For the TBAJ-876 dataset, states 1, 2, and 3 were calculated from 96,742, 58,915, and 166,169 particles images and reached resolutions of 2.8, 2.9, and 2.6 Å, respectively. For the SQ31f dataset, states 1, 2, and 3 were calculated from 101,399, 47,332 and 79,965 particles images and reached resolutions of 2.8, 3.1, and 2.9 Å, respectively. Masked local refinement of the F_O_ region yielded a map at 2.8 Å resolution for the TBAJ-876 dataset, and a map at 2.8 Å resolution for the SQ31f dataset.

### Atomic model building and refinement

The BDQ-bound ATP synthase models were fit as rigid bodies into the SQ31f and TBAJ-876 maps with UCSF Chimera (Goddard et al., 2007). Adjustments to the models were made in Coot (Emsley and Cowtan, 2004). ISOLDE (Croll, 2018) and PHENIX (Afonine et al., 2018) were then used to refine the models. Model quality was evaluated with Molprobity (Chen et al., 2010) and EMRinger (Barad et al., 2015). Figures and movies were generated with UCSF Chimera and ChimeraX (Goddard et al., 2018).

## Data availability

All maps and models are available through the Electron Microscopy Databank through accession codes EMD-29648, EMD-29649, EMD-29650, EMD-29651, EMD-29652, EMD-29653, EMD-29654, EMD-29655, and Protein Databank with accession codes 8G07, 8G08, 8G09, 8G0A, 8G0B, 8G0C, 8G0D, 8G0E.

## Acknowledgements

We thank Dr. Christopher Cooper and TB Alliance for providing TBAJ-587 and TBAJ-876. GMC was supported by an Ontario Graduate Scholarship for International Students and JLR was supported by the Canada Research Chairs program. This research was supported by CIHR grant PJT162186 (JLR) and Deutsche Forschungsgemeinschaft (DFG, German Research Foundation) grant 432291016 (AR). CryoEM data were collected at the Toronto High-Resolution High-Throughput Cryo-EM facility, supported by the Canada Foundation for Innovation and Ontario Research Fund.

## Statement of contributions

GMC performed enzyme assays and structural analysis. PP synthesized SQ31f. LM measured MIC and MBC for SQ31f with *M. smegmatis*. AR supervised the measurement of MIC and MBC. PI supervised synthesis of SQ31f and coordinated experiments. JLR conceived the study and coordinated experiments. GMC and JLR wrote the manuscript and prepared the figures with input from the other authors.

## Supplementary Figures

**Supplementary Figure 1.**
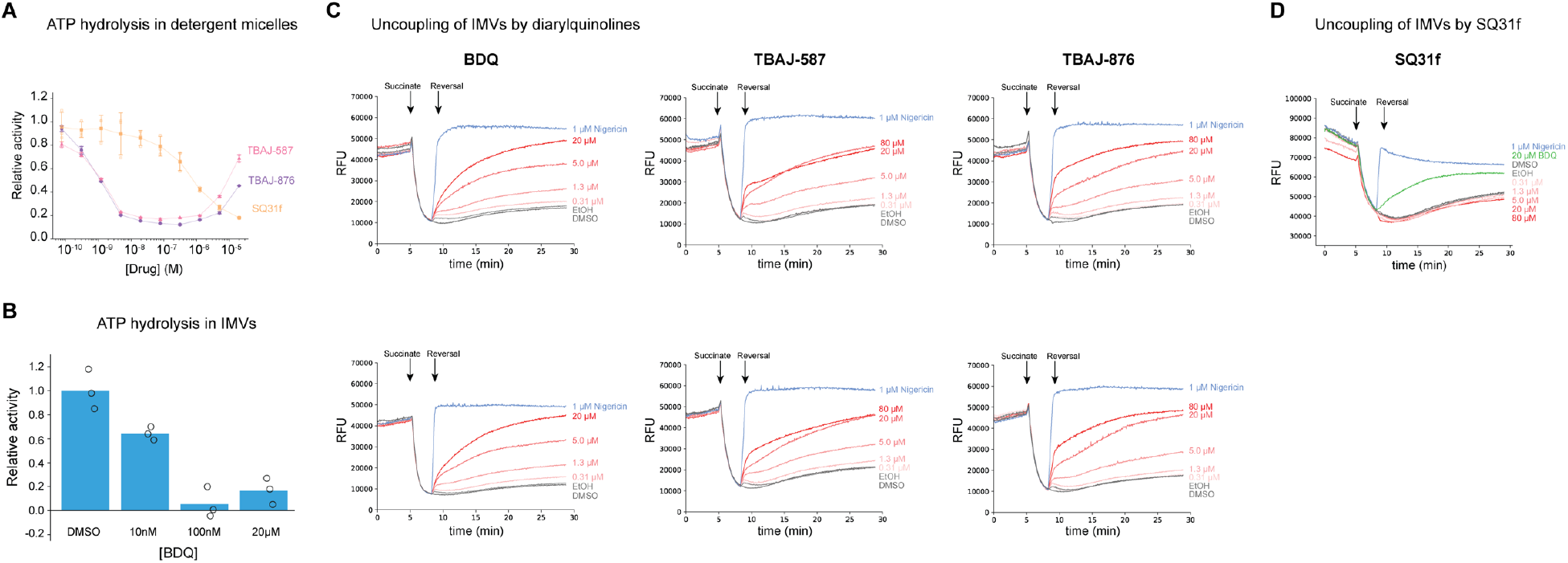
Replicate data for ATPase inhibition and IMV uncoupling by diarylquinolines and SQ31f. **A,** Inhibition of ATP hydrolysis by diarylquinoline analogs and SQ31f with purified ATP synthase bearing truncated α subunits. Mean relative activity (filled symbols) ± s.d from n=3 separate assays (empty symbols) for TBAJ-876, TBAJ-587, and SQ31f (triangles, circles, and squares). **B,** Inhibition of ATP hydrolysis activity in *M. smegmatis* IMVs containing ATP synthase with truncated α subunits. ATP hydrolysis activity by the ATP synthase is defined as the hydrolytic activity that is sensitive to 100 μM DCCD. **C,** Uncoupling activity of diarylquinoline analogs in *M. smegmatis* IMVs monitored with ACMA. Titration curves are shown for two independent experiments for each of three different inhibitors. **D,** Uncoupling activity of SQ31f in *M. smegmatis* IMVs monitored with acridine orange.

**Supplementary Figure 2.**
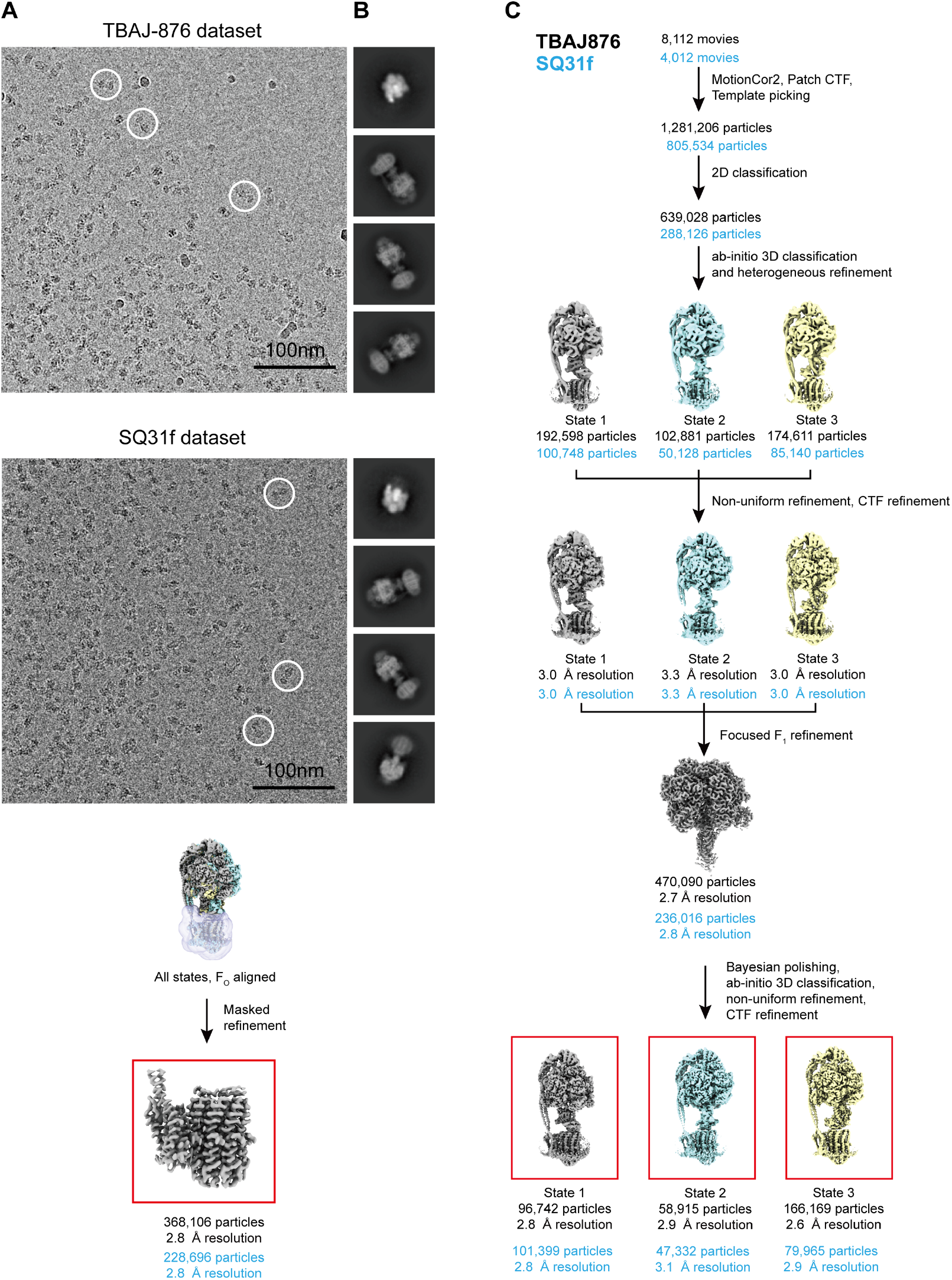
Workflow for Cryo-EM image analysis of TBAJ-876 and SQ31f bound *M. smegmatis* ATP synthase. **A,** Example micrograph from the TBAJ-876 dataset (upper) and the SQ31f dataset (lower) with representative ATP synthase particles circled. **B,** 2D class average images from the datasets. **C,** Workflow followed to obtain maps of the three rotational states and focused refinement of the F_O_ region with TBAJ-876 (black particle image counts and resolutions) and SQ31f (blue particle image counts and resolutions). 3D maps shown to illustrate the workflow are from the TBAJ-876 dataset.

**Supplementary Figure 3.**
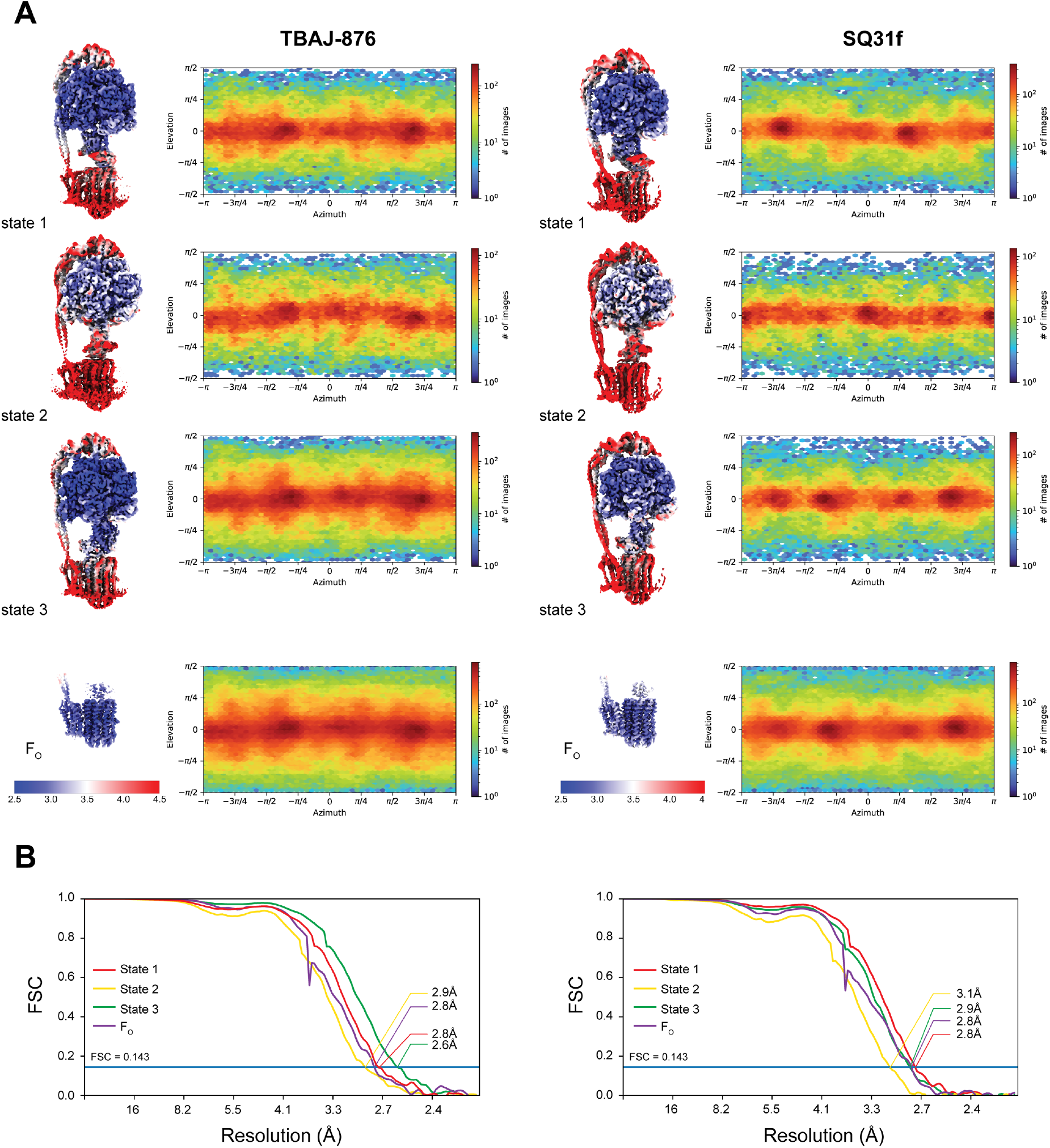
Cryo-EM map validation. **A,** Local resolution and particle orientation distribution plots are shown for maps of the three rotational states and the *F*_O_ regions with TBAJ-876 and SQ31f bound. **B,** Fourier shell correlation curves corrected for masking following a gold-standard refinement are shown for all maps.

**Supplementary Figure 4.**
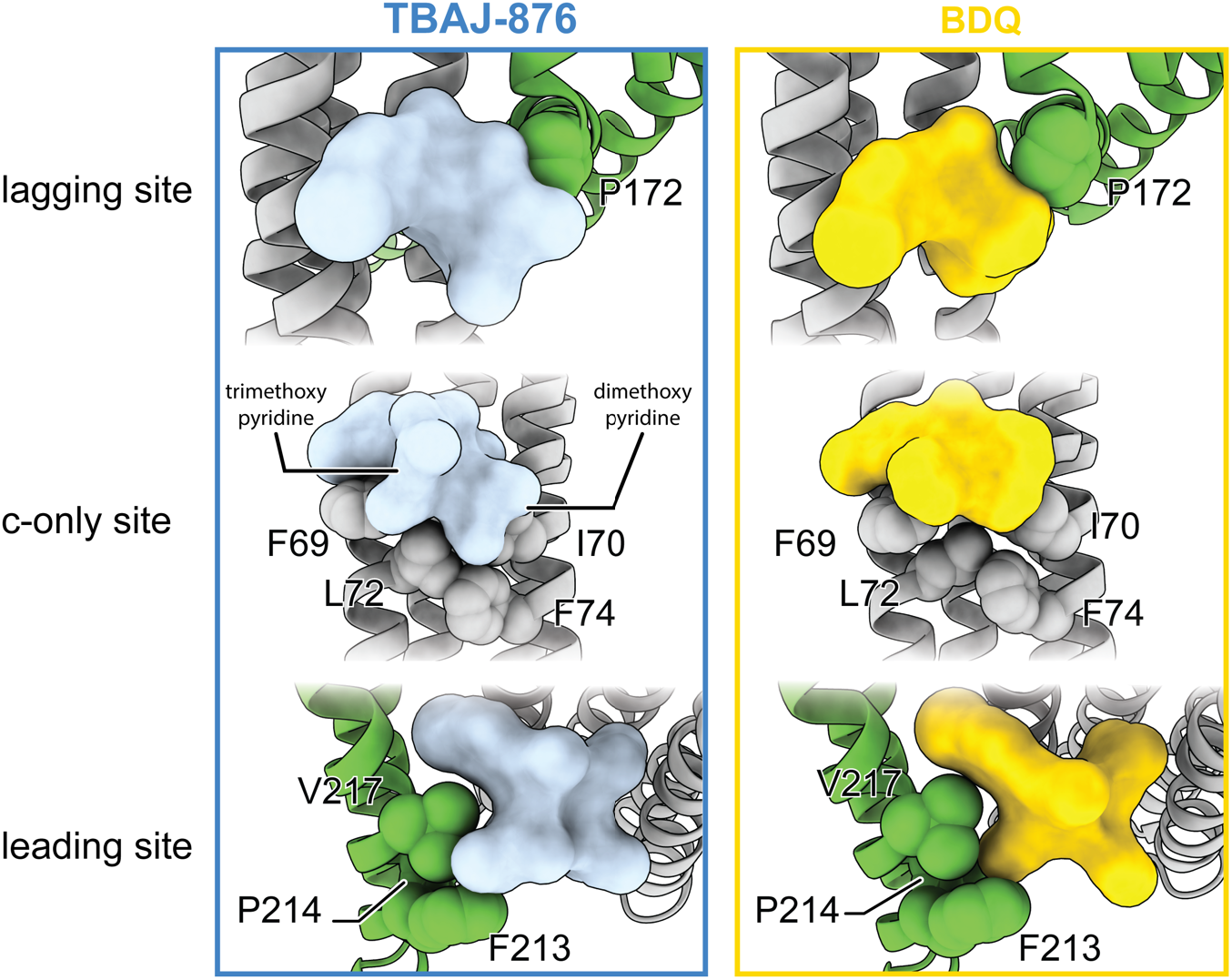
Comparison of BDQ and TBAJ-876 binding sites. Atomic models of BDQ (right, from (Guo et al., 2021)) and TBAJ-876 (left) in the lagging, c-only, and leading site of *M. smegmatis* ATP synthase. TBAJ-876 forms improved interactions compared to BDQ with specific residues, which are displayed as space filling models.

**Supplementary Figure 5.**
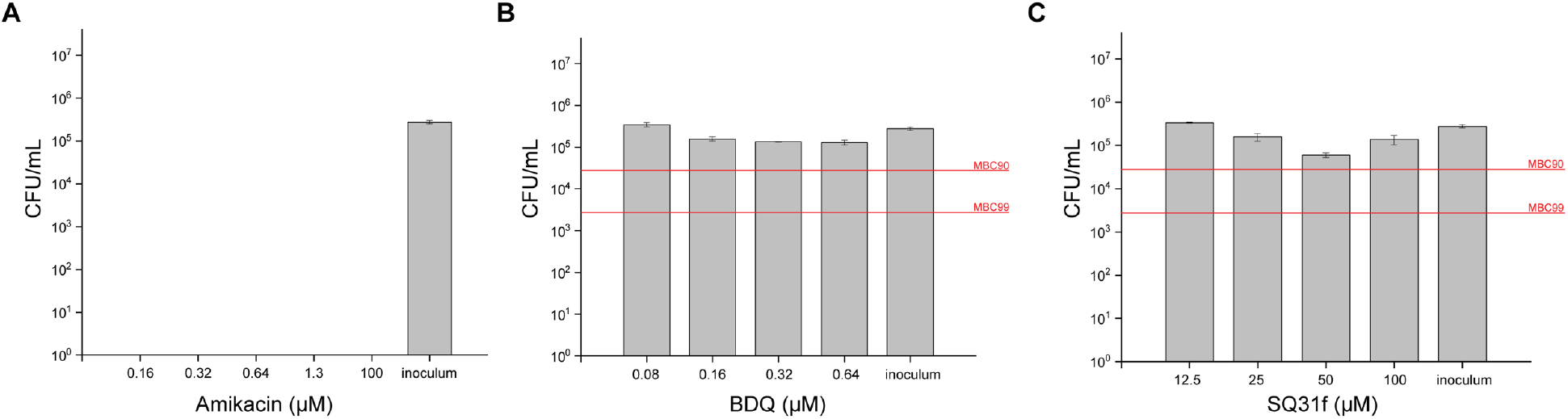
Minimum bactericidal concentration assays for BDQ and SQ31f. Minimal bactericidal concentration assay for Amikacin (**A**), BDQ (**B**), and SQ31f (**C**). BDQ and SQ31f bactericidal activity was assayed at up to 8 × MIC_90_. Thresholds for MBC90 and MBC99 are shown as red lines.

**Supplementary Table 1.** Cryo-EM data collection, refinement, and validation statistics.

|  | **TBAJ876 state 1**<br>(EMD-29653,<br>PDB 8G0C) | **TBAJ876 state2**<br>(EMD-29654,<br>PDB 8G0D) | **TBAJ876 state3**<br>(EMD-29655,<br>PDB 8G0E) | **TBAJ876 focused F<sub>0</sub>**<br>(EMD-29652,<br>PDB 8G0B) |
| --- | --- | --- | --- | --- |
| Magnification | 75,000 | 75,000 | 75,000 | 75,000 |
| Voltage (kV) | 300 | 300 | 300 | 300 |
| Electron exposure (e<sup>-</sup>/Å<sup>2</sup>) | 44 | 44 | 44 | 44 |
| Exposure rate (e<sup>-</sup>/pixel/s) | 7 | 7 | 7 | 7 |
| Exposure time (s) | 6.7 | 6.7 | 6.7 | 6.7 |
| Defocus range (μm) | 0.7-2.1 | 0.7-2.1 | 0.7-2.1 | 0.7-2.1 |
| Pixel size (Å) | 1.03 | 1.03 | 1.03 | 1.03 |
| Symmetry imposed | C1 | C1 | C1 | C1 |
| Initial particle images (no.) | 1,281,206 | 1,281,206 | 1,281,206 | 1,281,206 |
| Final particle images (no.) | 96,742 | 58,915 | 166,169 | 368,106 |
| Map resolution (Å) | 2.8 | 2.9 | 2.6 | 2.8 |
| FSC threshold | 0.143 | 0.143 | 0.143 | 0.143 |
| Map resolution range (Å) | 2.3-5.4 | 2.5-6.8 | 2.3-5.5 | 2.3-3.4 |
| **Refinement** |  |  |  |  |
| Initial model used (PDB code) | 7JGA | 7JGA | 7JGA | 7JGC |
| Model resolution (Å) |  |  | 3.3 | 2.9 |
| FSC threshold |  |  | 0.5 | 0.5 |
| Map sharpening B factor (Å<sup>2</sup>) |  |  | locally sharpened | Locally sharpened |
| **Model composition** |  |  |  |  |
| Non-hydrogen atoms |  |  | 35,959 | 7,758 |
| Protein residues |  |  | 4858 | 1044 |
| Ligands |  |  | 4 mgATP<br>1 PO4<br>7 TBAJ876 | 7 TBAJ876 |
| **R.m.s deviations** |  |  |  |  |
| Bond lengths (Å) |  |  | 0.009 | 0.007 |
| Bond angles (°) |  |  | 1.156 | 0.961 |
| **Validation** |  |  |  |  |
| MolProbity score |  |  | 1.55 | 1.58 |
| Clashscore |  |  | 7.06 | 7.57 |
| Poor rotamers (%) |  |  | 1.21 | 0.00 |
| Favored (%) |  |  | 97.52 | 97.05 |
| Allowed (%) |  |  | 2.44 | 2.95 |
| Disallowed (%) |  |  | 0.04 | 0.00 |

|  | **SQ31f state 1**<br>(EMD-29649,<br>PDB 8G08) | **SQ31f state2**<br>(EMD-29650,<br>PDB 8G09) | **SQ31f state3**<br>(EMD-29651,<br>PDB 8G0A) | **SQ31f focused F<sub>0</sub>**<br>(EMD-29648,<br>PDB 8G07) |
| --- | --- | --- | --- | --- |
| Magnification | 75,000 | 75,000 | 75,000 | 75,000 |
| Voltage (kV) | 300 | 300 | 300 | 300 |
| Electron exposure (e<sup>-</sup>/Å<sup>2</sup>) | 54 | 54 | 54 | 54 |
| Exposure rate (e<sup>-</sup>/pixel/s) | 6.9 | 6.9 | 6.9 | 6.9 |
| Exposure time (s) | 8.3 | 8.3 | 8.3 | 8.3 |
| Defocus range (μm) | 0.9-2.0 | 0.9-2.0 | 0.9-2.0 | 0.9-2.0 |
| Pixel size (Å) | 1.03 | 1.03 | 1.03 | 1.03 |
| Symmetry imposed | C1 | C1 | C1 | C1 |
| Initial particle images (no.) | 805,534 | 805,534 | 805,534 | 805,534 |
| Final particle images (no.) | 101,399 | 47,332 | 79,965 | 228,696 |
| Map resolution (Å) | 2.8 | 3.1 | 2.9 | 2.8 |
| FSC threshold | 0.143 | 0.143 | 0.143 | 0.143 |
| Map resolution range (Å) | 2.3-5.5 | 2.7-6.4 | 2.4-5.6 | 2.4-3.5 |

| Refinement |  |  |  |  |
| --- | --- | --- | --- | --- |
| Initial model used (PDB code) | 7JGA | 7JGA | 7JGA | 7JGC |
| Model resolution (Å) |  |  | 3.3 | 2.9 |
| FSC threshold |  |  | 0.5 | 0.5 |
| Map sharpening *B* factor (Å<sup>2</sup>) |  |  | locally sharpened | Locally sharpened |
| Model composition |  |  |  |  |
| Non-hydrogen atoms |  |  | 35,641 | 7,467 |
| Protein residues |  |  | 4861 | 1045 |
| Ligands |  |  | 4 mgATP<br>1 PO4<br>1 SQ31f | 1 SQ31f |
| R.m.s deviations |  |  |  |  |
| Bond lengths (Å) |  |  | 0.007 | 0.008 |
| Bond angles (°) |  |  | 1.082 | 1.061 |
| Validation |  |  |  |  |
| MolProbity score |  |  | 1.53 | 1.55 |
| Clashscore |  |  | 7.64 | 7.46 |
| Poor rotamers (%) |  |  | 1.12 | 1.15 |
| Favored (%) |  |  | 97.65 | 97.55 |
| Allowed (%) |  |  | 2.31 | 2.45 |
| Disallowed (%) |  |  | 0.04 | 0.00 |

**Video 1.** Binding of BDQ (yellow), TBAJ-876 (blue), and SQ31f (red) to the mycobacterial ATP synthase. The movie shows interpolation between the drug-free and drug-bound conformations of the mycobacterial ATP synthase in rotational state 1.

